## Supplemental Figures for "Cryo-electron tomography of Birbeck granules reveals the molecular mechanism of langerin lattice formation"

### Supplemental Materials

**Supplemental Figure 1-3**

**Supplemental Movie 1 and 2**

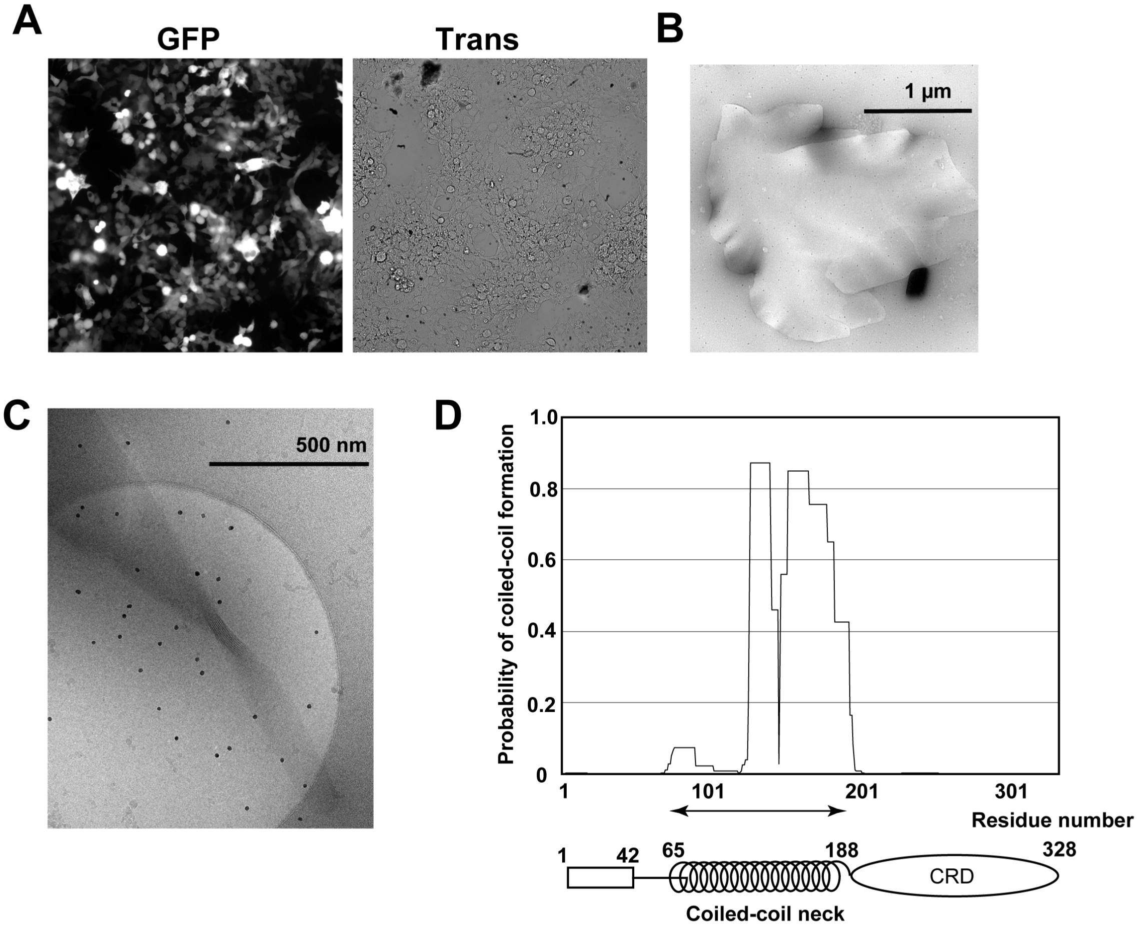


### Supplemental Figure 1

**Supplemental Figure 1: Isolation of Birbeck granules. Related to Figure 1.** (A) Fluorescence microscopy image of 293T cells expressing langerin. IRES-driven GFP expression was detected. The transformation efficiency was approximately 40%. (B) Negative-stain electron microscopy image of an isolated Birbeck granule stained with 2% uranium acetate. (C) Cryo-electron microscopy image of a twisted Birbeck granule. (D) Coiled-coil prediction profile of langerin. (E) Effect of expression level in Birbeck granule formation. Immunoblots showed low langerin expression in the stable cell line. Due to the low langerin expression, Birbeck granule formation was also low.

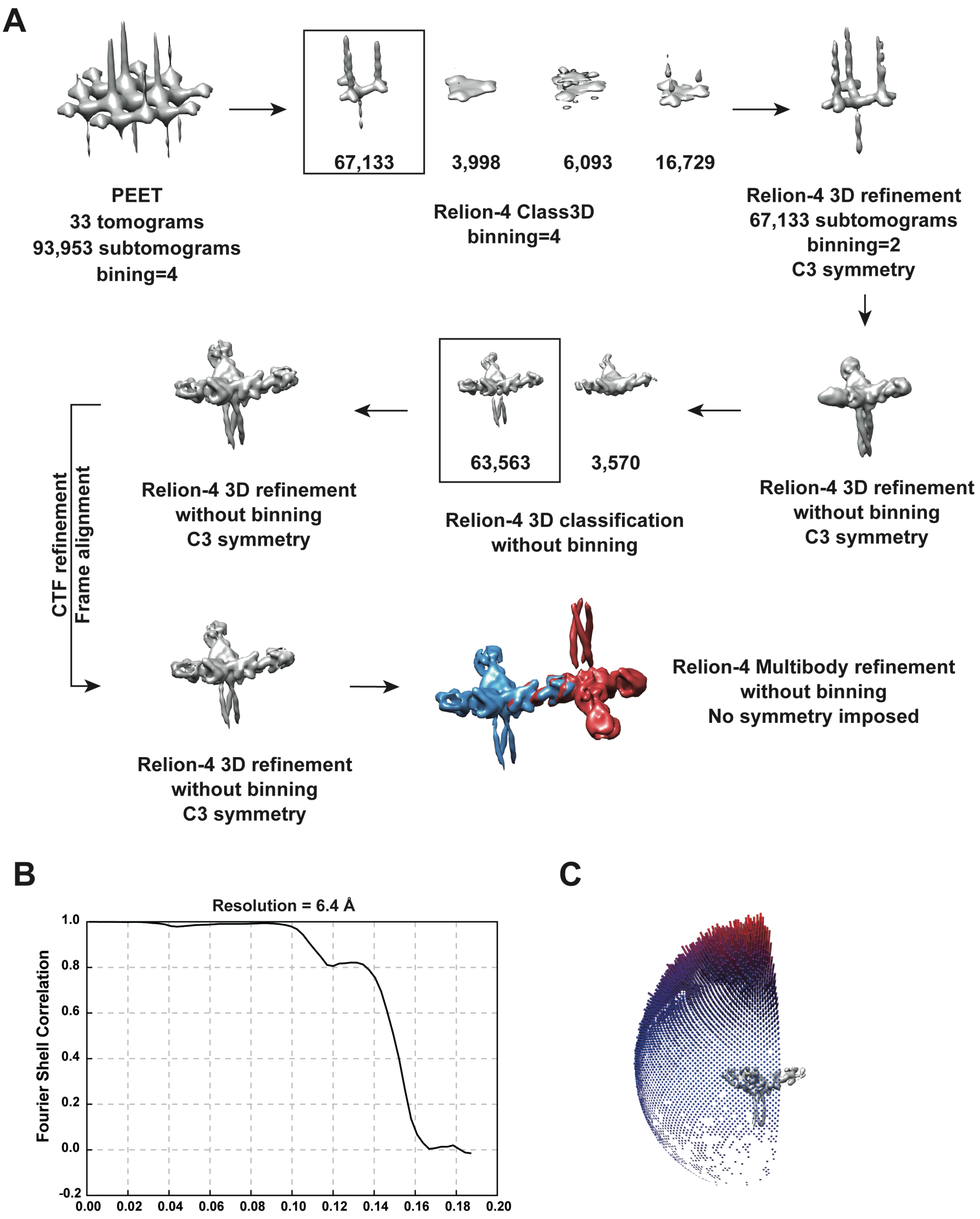


**Supplemental Figure 2**

**Supplemental Figure 2: Cryo-electron tomography of Birbeck granules. Related to**

**Figure 2.** (A) Summary of image processing. (B) Fourier shell correlation plot of the 3D refined structure of the langerin trimer, corresponding to the map in Figure 2C. The estimated resolution was 6.4 Å with a cut-off value of 0.143. (C) Angular distribution of the subtomograms. Although there was a strong bias toward the top-view direction, twisting of some granules (Figure S1D) provided side views.

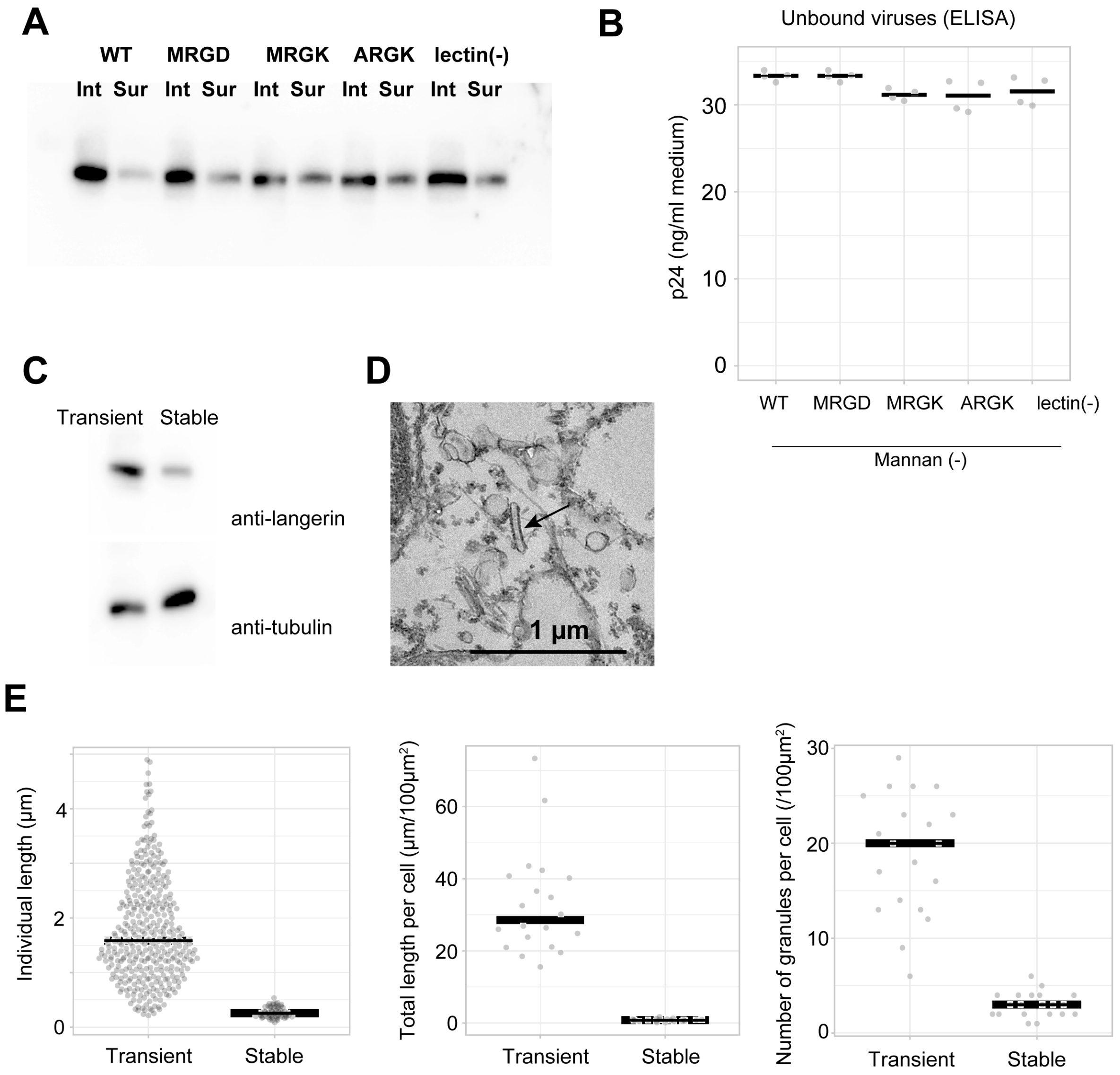


**Supplemental Figure 3**

**Supplemental Figure 3: Virus internalization by langerin. Related to Figure 4 and 5.**

(A) Electron microscopy of HIV-infected cells. 293T cells expressing wild type langerin were incubated with HIV pseudovirus for 10 min at 37℃, and the internalized viruses (arrows) were visualized by ultra-thin section electron microscopy. (B) Surface labeling of langerin.

Langerin-expressing cells were surface-labeled using biotin N-hydroxysulfosuccinimide ester, and labeled langerin were immunoprecipitated using streptavidin agarose. “Int” and “Sur” indicate intracellular and surface langerin, respectively. The intracellular : surface ratio of langerin was approximately 5:1 and this ratio was not significantly affected by mutations. (C) Quantification of unbound viruses using p24 ELISA. Medium supernatants were diluted 1000-fold before loading into the ELISA plate. Concentration of viruses in the medium supernatant were nearly equal among the wild type and the mutants. (D) Anti-langerin immunoblot of stably-expressing cell line. The expression level of langerin in the stable cell line was approximately 30% of that of transiently-expressing cells. Given that the transformation efficiency of transiently-expressing cells was 40%, the actual expression level of langerin per cell is estimated to be ~12%. (E) Electron microscopy of the stable cell line. Birbeck granule formation was induced by addition of yeast mannan. Short and isolated Birbeck granules were observed (arrow) (F) Quantification of Birbeck granule formation. “Transient” corresponds to WT mannan (+) in Figure 4B-D. 74 Birbeck granules in 20 stably-expressing cells were measured.

**Supplemental Movie 1: Related to Figure 2. Movie representation of the langerin lattice.**

**Supplemental Movie 2: Related to Figure 3B. Morphing movie** showing the first eigenvector of structural variation calculated by multibody analysis.

**Table S1: Summary of data collection and model validation**

| **Data collection parameters** | |
|---|---|
| Magnification | 33,000× |
| Pixel size (Å) | 2.67 |
| Defocus range (µm) | 2.6 – 8.5 |
| Voltage (keV) | 300 |
| Exposure time (sec/frame) | 0.74 |
| Number of frames per tilt | 20 |
| Angular range (°) | ±60 |
| Increments (°) | 3 |
| Total dose ($e^-$/Å$^2$) | 49.6 |
| Box size (pixels) | 128 |
| Number of tilt series recorded | 126 |
| Number of tilt series processed | 33 |
| Initial number of subtomograms | 93,953 |
| Final number of subtomograms | 63,563 |
| **Model validation statistics** | |
| Initial model used (PDB ID) | 3KQG |
| Bonds length RSMD (Å) | 0.003 |
| Bonds angles RSMD (°) | 0.651 |
| MolProbability score | 1.83 |
| Clash score | 21.13 |
| Rotamer outliers (%) | 0.0 |
| Ramachandran plot (%) | Favored: 97.94<br>Allowed: 2.06<br>Outliers: 0.00 |
| CaBLAM outliers (%) | 1.14 |
| B-factors (min/max/mean) | 443.64/1003.68/628.65 |
| Map resolution estimates (Å) | 6.4 ($FSC_{half\text{-}map}$=0.143)<br>8.5 ($FSC_{model}$=0.143) |
